# Hillslope geodiversity shapes ammonia-oxidizing communities and other microbial regulators in a semi-arid shrubland

**DOI:** 10.1101/2021.03.08.434393

**Authors:** Amir Szitenberg, Rivka Alexander-Shani, Hezi Yizhak, Ilan Stavi

## Abstract

The determinants and consequences of drought-related shrub mortality were studied for over a decade, as a model for desertification processes, in a semi-arid long-term ecological research station. Recent studies have shown that geodiversity is an important spatial predictor of plant viability under extreme drought conditions. Homogeneous hillslopes, with a deep soil profile and lack of stoniness, could not support shrubs under long term drought conditions due to low water storage in their soil. Conversely, heterogeneous hillslopes, with shallow soil profiles and high stoniness, supported shrub communities under similar conditions, due to the comparatively greater soil-water content. In the current study, we investigated the effect of hillslope geodiversity on the soil microbial diversity. Using DNA metabarcoding, we found small but consistent differences in the microbial community compositions of the homogeneous and heterogeneous hillslopes; more ammonia oxidizing and reducing-sugar degrading bacteria are found in the homogeneous hillslopes, possibly dwindling the ammonia supply to shrubs. Additionally, based on functional metagenomic reconstruction, we suggest that homogeneous hillslopes have lower superoxide and antibiotics production, leading to reduced protection against pathogens. In fungi, we observed an increase in possible pathogens, at the expense of lichen forming fungi. Lichens are considered to support soil-water by slowly releasing intercepted raindrops. In conclusion, we show that not only plant-diversity but also microbial-diversity is shaped by geodiversity, and that the community shift in homogeneous hillslopes may further promote shrub mortality in this drought-prone, water limited ecosystem.

**HIGHLIGHTS:**

- Homogeneous hillslopes reduce soil water storage and increase aeration.
- Ammonia oxidizers and reducing-sugar degraders dwindle ammonia supply for plants.
- Homogenous hillslopes do not support moisture providing lichens.
- Reduced antibiotics and superoxide secretion capacitate pathogens.
- Geodiversity facilitates microbial regulation during drought.

**GRAPHICAL ABSTRACT:** 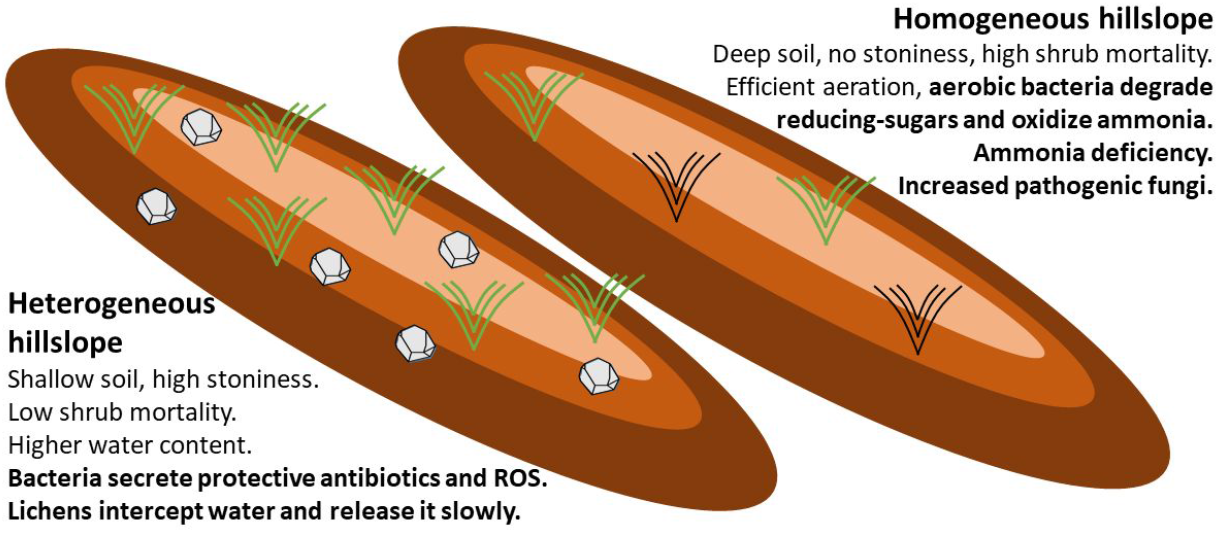

## 1. Introduction

Climate change is predicted to alter precipitation regimes toward more frequent and severe drought events (Shukla et al., 2019). Consequently, understanding the direct effects of long term shifts in rainfall quantities and frequencies on biodiversity has been a major focus of ecological research (Beier et al., 2012; Hoover et al., 2014; Kreyling and Beier, 2013) and a key challenge for land managers (Briske et al., 2015; Smith, 2011). Such shifts may cause ecosystem transitions, which may lead to considerable and irreversible changes in biodiversity and ecosystem services (Díaz et al., 2019). Soil moisture changes directly affect plant productivity and survival, as well as mobilization of soil minerals and the diversity and activity of soil and plant microbiota (Dijkstra et al., 2015). The soil microbiome, in turn, plays vital roles in the cycling, mobilization and transport of nutrients, plant growth promotion, biocontrol of phytopathogens, water uptake capacity and abiotic stress response (Sze et al., 2020).

The spatial differences in lithological, topographic, and pedological properties of the soil, commonly regarded as geodiversity (Gray, 2005), are known to locally alter water storage in the soil profile and the mobilization of minerals (Okin et al., 2015; Stavi et al., 2018a). The interaction between this physical background and rainfall patterns, particularly in water-limited ecosystems, modulates the effects of climate change on biodiversity. Geodiversity effects on biodiversity have been studied mostly for plants and animals (Cartwright, 2019; Ibáñez and Feoli, 2013; Read et al., 2020; Stavi et al., 2018b; Stein et al., 2014), and for large geographic scales, such as watersheds and landscapes (Ibáñez et al., 2012; Janion-Scheepers et al., 2016). Millimetric scale pedodiversity as a driver of microbial diversity has also been investigated (Upton et al., 2019). Here, we aim to study the effects of geodiversity on the soil microbial diversity at the hillslope scale.

The Sayeret-Shaked Park in the Negev region of southern Israel is an established study site of regional desertification processes (Sher et al., 2012). Small-scale geodiversity in the region has been generated by the uneven spatial distribution of livestock traffic on the hillslopes (Stavi et al., 2018b, 2015). Concomitantly, hillslope-scale geodiversity was attributed to stoniness and depth of the soil profile (Stavi et al., 2019; Yizhaq et al., 2017). In both cases, elevated geodiversity was shown to alleviate the shortage of water for shrubs, increasing their durability under decreased precipitation regimes (Stavi et al., 2019; Yizhaq et al., 2017). Localized patches of high geodiversity were found to be critical for the survival of *Noaea mucronata* (Forssk.) Asch. & Schweinf, the most common shrub species across the study area, under prolonged drought (Stavi et al., 2019). In this study we set out to understand the impact of geodiversity variation, generated by differences in stoniness and soil depth, on the bacterial and fungal communities in the soil and the consequent microbial functions. We hypothesized that the microbial community composition would differ between the hillslope types, because many microbes respond rapidly to changes in abiotic condition, with consequences to regulatory ecosystem services.

## Materials and methods

### 2.1 The study site

The study was carried out in the Long Term Ecological Research (LTER) station of the Sayeret Shaked Park (31°27’ N, 34°65’ E; 187 m.a.s.l.), in the semi-arid north-western Negev of southern Israel. The station, which covers an area of ca. 20 ha, has been fenced since the 1990s to prevent livestock grazing. The lithology is Eocene chalk and Plio-Pleistocene eolianites. The landform is dominated by rolling hills, with hillslope incline ranging between 3° and 6°. The soil is classified as loessial Calcic Xerosol, with a sandy loam to loamy sand texture (Stavi et al., 2018a). Depth and stoniness of soil, as well as rock fragment cover percentage, are highly varied in the region (Singer, 2007). Multiannual mean daily temperatures are 26 and 12°C in the summer and winter, respectively, and mean cumulative annual precipitation has been ~250mm (Bitan and Rubin, 1991). However, during the first decade of the twenty-first century, precipitation rates sharply decreased (Shachak, 2011), averaging only ~165mm·y^-1^. (Israel Meteorological Service website: http://www.ims.gov.il/ims/all_tahazit/). The reduced precipitation was followed by a trend of mass shrub mortality, as observed by Sher et al. (2012), particularly among *N. mucronata.* For further regional and floral context see Stavi et al. (2018a).

### 2.2 Sampling

During February 2020, at the peak of the growth season, samples were collected from six 20 × 20 m plots, located on six adjacent hillslopes (Fig. 1A). Three hillslopes, previously characterized as homogeneous, have a soil profile deeper than 1m and no stoniness, and three additional hillslopes, previously characterized as heterogeneous, have ~0.1 m soil layer and high stoniness, accounting for ~30% of the volume and ~35% cover of the ground surface (Stavi et al., 2018a). The minimal distance between each pair of plots was 100 m. To negate aspect-related and incline-related effects, all plots were delineated at a relatively similar azimuth (310° ± 29°) and slope (5° ± 0.5°). In each plot, we collected five soil samples from underneath the canopy of *N. mucronata* plants (shrubby patch) and five samples from an exposed area between shrubby patches (inter-shrub spaces), to reach a total of 60 soil samples. Each shrubby-patch sample was collected from an independent cluster of shrubs. To collect the samples, the soil was exposed with a trowel and a sterile 15 ml tube was inserted to the exposed soil at 8-10 cm depth. Samples were stored at −80°C until further processing.

**Fig. 1:**
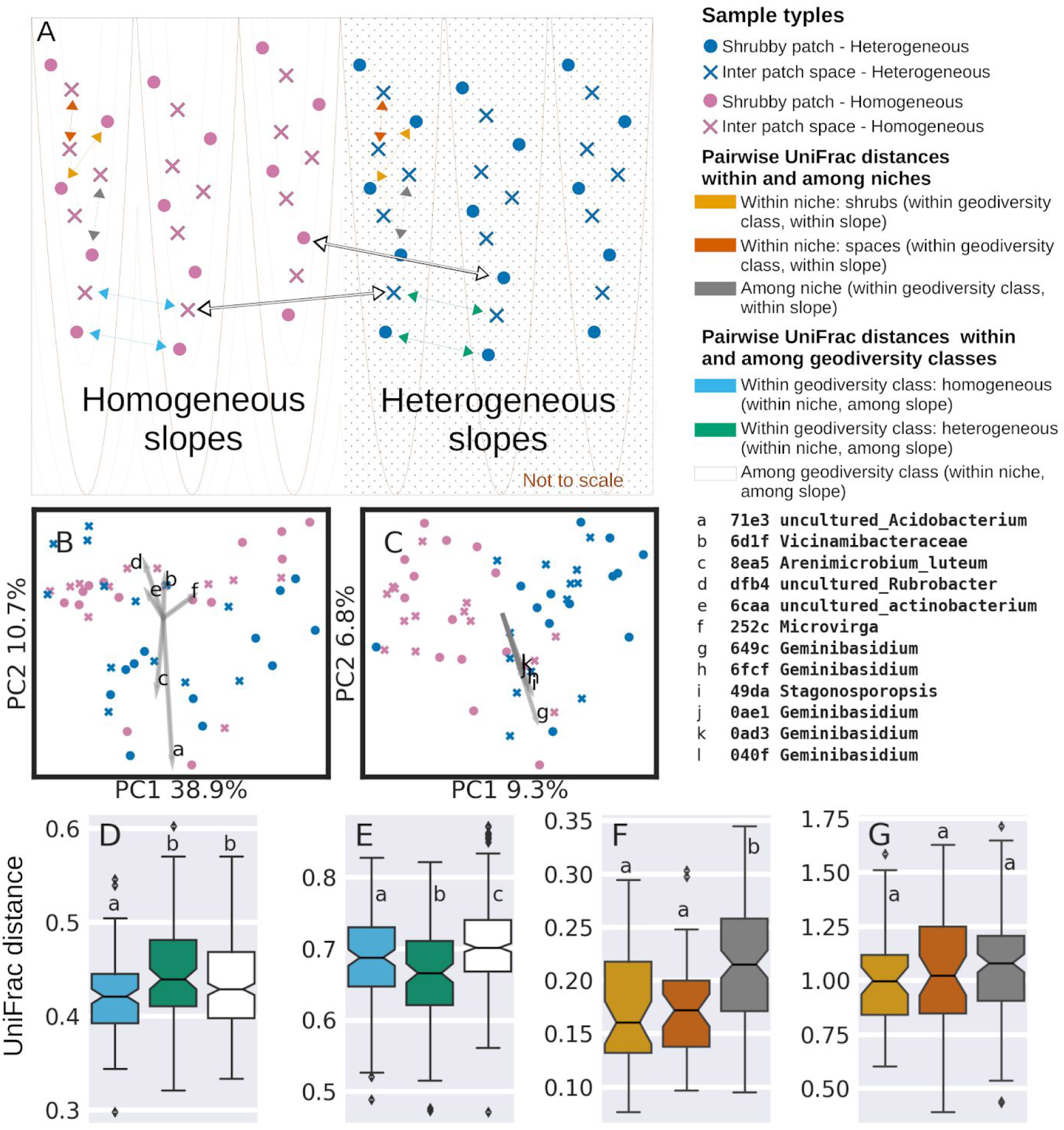
Investigation of microbial beta-diversity in the study area. Three homogeneous and three heterogeneous hillslopes were selected for this study. In each hillslope, five soil samples were collected from shrubby patches and five from inter-shrub areas (A), all from 8-10 cm depth. Arrows in sub-plot A exemplify the type of sample pairs which were used to calculate pairwise UniFrac distances for the beta diversity factorial ANOVA tests. The sample types are listed in the legend. Beta diversity was first investigated with UniFrac distance matrices and principal coordinate analyses (PCoA). Weighted analysis is presented for bacteria (B) and unweighted for fungi (C). Gray arrows (B and C) represent the six most important ASVs explaining the total variance, following the importance definition of Legendre and Legendre (2012). The identity of the important ASVs are indexed with lowercase letters and listed on the right-hand side. Unweighted UniFrac pairwise distances within and among biodiversity classes are presented as boxplots for prokaryotes (D) and fungi (E). Weighted UniFrac pairwise distances within and among niches are presented as boxplots for bacteria (F) and fungi (G). Within each boxplot (D-G), boxes which share a lowercase letter label represent distance distributions which are not significantly different, whereas boxes denoted with different letters represent distance distributions that are significantly different from each other.

### 2.3 16S rRNA and ITS1 metabarcoding

DNA was extracted using the DNeasy PowerSoil kit (Qiagen) following the manufacturer’s instructions. Metabarcoding libraries were prepared with a two step PCR protocol. For the first PCR reaction, the V4 16S rRNA region was amplified, following Toju et al. (2019), with the forward primer 515f 5’-tcg tcg gca gcg tca gat gtg tat aag aga cag GGT GCC AGC MGC CGC GGT AA-3’ and the reverse primer 806R 5’-gtc tcg tgg gct cgg aga tgt gta taa gag aca gGA CTA CHV GGG TWT CTA AT-3’, along with artificial overhang sequences (lowercase). For ITS1, the first PCR primers were ITS1_F_KYO2 5’-TAG AGG AAG TAA AAG TCG TAA-3’ and ITS2_KYO1 5’-gtc tcg tgg gct cgg aga tgt gta taa gag aca gCT RYG TTC TTC ATC GDT-3’ following Toju et al. (2012), with the same overhangs as in the V4 primer. In the second PCR reaction, sample specific barcode sequences and Illumina flow cell adapters were attached, using the forward primer ‘5-AAT GAT ACG GCG ACC ACC GAG ATC TAC ACt cgt cgg cag cgt cag atg tgt ata aga gac ag-’3 and the reverse primer ‘5-CAA GCA GAA GAC GGC ATA CGA GAT XXX XXX XXg tct cgt ggg ctc gg-’3’, including Illumina adapters (uppercase), overhang complementary sequences (lowercase), and sample specific DNA barcodes (‘X’ sequence). The PCR reactions were carried out in triplicate, with the KAPA HiFi HotStart ReadyMix PCR Kit (KAPA biosystems), in a volume of 25 μl, including 2 μl of DNA template and following the manufacturer’s instructions. The first PCR reaction started with a denaturation step of 3 minutes at 95 °C, followed by 30 cycles of 20 seconds denaturation at 98°C, 15 seconds of annealing at 55°C and 7 seconds polymerization at 72°C. The reaction was finalized with another minute long polymerization step. The second PCR reaction was carried out in a volume of 25 μl as well, with 2 μl of the PCR1 product as DNA template. It started with a 3-minutes denaturation step at 95°C, followed by 8 cycles of 20 seconds denaturation at 98°C, 15 seconds of annealing at 55°C and 7 seconds polymerization at 72°C. The second PCR reaction was also finalized with another 60-second polymerization step. The first and second PCR reaction products were purified using AMPure XP PCR product cleanup and size selection kit (Beckman Coulter), following the manufacturer’s instructions, and normalised based on Quant-iT PicoGreen (Invitrogen) quantifications. The fragment size distribution in the pooled libraries was examined on a TapeStation 4200 (Agilent) and the libraries were sequenced on an iSeq-100 Illumina platform, producing 150 bp paired end reads. Sequence data was deposited in the National Center for Biotechnology Information (NCBI) data bank, under BioProject accession PRJNA697940.

### 2.4 Data processing, taxonomy assignment and biodiversity analysis

All the analysis carried out for this study is available as a Jupyter notebook in a github repository (GitHub: https://git.io/Jt4WP, Zenodo DOI: 10.5281/zenodo.4480328), along with the sequence data, intermediate and output files. The bioinformatics analysis was carried out with Qiime2 (Bolyen et al., 2019). DADA2 (Callahan et al., 2016) was used to trim PCR primers, quality-filter, error correct, dereplicate and merge the read pairs, and to remove chimeric sequences, to produce the amplicon sequence variants (ASV). Specific ASVs are referred to by the first four characters of their MD5 digests, which correspond with the biom table headers. For taxonomic assignment, a naive Bayes classifier was trained using taxonomically identified reference sequences from the Silva 138 SSU-rRNA database (Quast et al., 2013) for the V4 fragments and the UNITE Feb 2019 database (Abarenkov et al., 2010) for the ITS1 fragments. All ASVs that were identified as mitochondrial or chloroplast sequences were filtered out from the feature table. The filtered ASV biom table was further filtered to exclude samples with less than 3000 sequences. These cutoffs were determined based on alpha rarefaction curves and represented the lowest read count in the saturated section of the curve. An ASV phylogenetic tree was built with the q2-phylogeny plugin, implementing MAFFT 7.3 (Katoh and Standley, 2013) for sequence alignment, and FastTree 2.1 (Price et al., 2010), with the default masking options. Microbial diversity was estimated based on Faith’s phylogenetic diversity (Faith, 1992) for alpha diversity and weighted and unweighted UniFrac distance (Lozupone and Knight, 2005) matrices for beta diversity. Ordination of the beta-diversity distance was carried out with a principal coordinates analysis (PCoA; Halko et al., 2011; Legendre and Legendre, 2012). The effects of the hillslope-type, niche and hillslope on the microbial diversity were tested with a factorial ANOVA using the q2-longitudinal plugin (Bokulich et al., 2017). P-values were corrected for multiple testing using the Benjamini-Hochberg procedure (Benjamini and Hochberg, 1995). Corrected p-values are referred to as q-values throughout the text.

### 2.5 Identification of explanatory taxa, ASVs and prokaryotic genes

To identify differentially abundant orders between the homogeneous and heterogeneous hillslopes we collapsed the ASV table to the order level and carried out pairwise Mann-Whitney tests (Mann and Whitney, 1947) between the two classes. As before, p-values were corrected for multiple testing using the Benjamini-Hochberg procedure (Benjamini and Hochberg, 1995). ASVs explaining the total variance of UniFrac distances were identified with a biplot analysis (Legendre and Legendre, 2012). ASV differences between the two hillslope types were investigated with a supervised learning approach based on logistic regression (Yu et al., 2011) using the Scikit-learn package (Pedregosa et al., 2011). Data were equally split between the training and testing sets, and were stratified to equally represent homogeneous and heterogeneous hillslopes in each subset. The procedure was replicated across 100 different random-seeds for the train-test data splitting step. Logistic regression was also used to identify explanatory relative abundance changes of prokaryotic genes, between the two geodiversity classes. 16S rRNA based metagenome reconstruction was carried out with PiCRUST2 (Langille et al., 2013).

## 3. Results

We obtained 55 samples with a median coverage of 7679 reads for the V4 fragment and 54 samples with a median coverage of 8847 reads for the ITS1 fragment, after the exclusion of low quality, chimeric and organelle sequences, and low coverage samples (< 3000 post-filtration read-pairs). The prokaryote and fungal orders with the highest average relative abundances are presented in Fig. 2. On average, the most abundant bacterial orders were Rubrobacterales (0.12 and 0.09 for heterogeneous and homogeneous hillslopes, respectively), Vicinamibacterales (0.1 and 0.096), Pyrinomonadales (0.09 and 0.06), and Rhizobiales (0.07 and 0.08). The most abundant fungal orders were, on average, Pleosporales (0.19 and 0.24), Sordariales (0.15 and 0.07), Mortierellales (0.076 and 0.079) and Xylariales (0.02 and 0.05). On the order level, relative abundances were similar between the two hillsploe types, except for the bacterial orders Micrococcales and Burkholderiales, and the fungal order Chaetothyriales (q-value < 0.05).

**Fig. 2:**
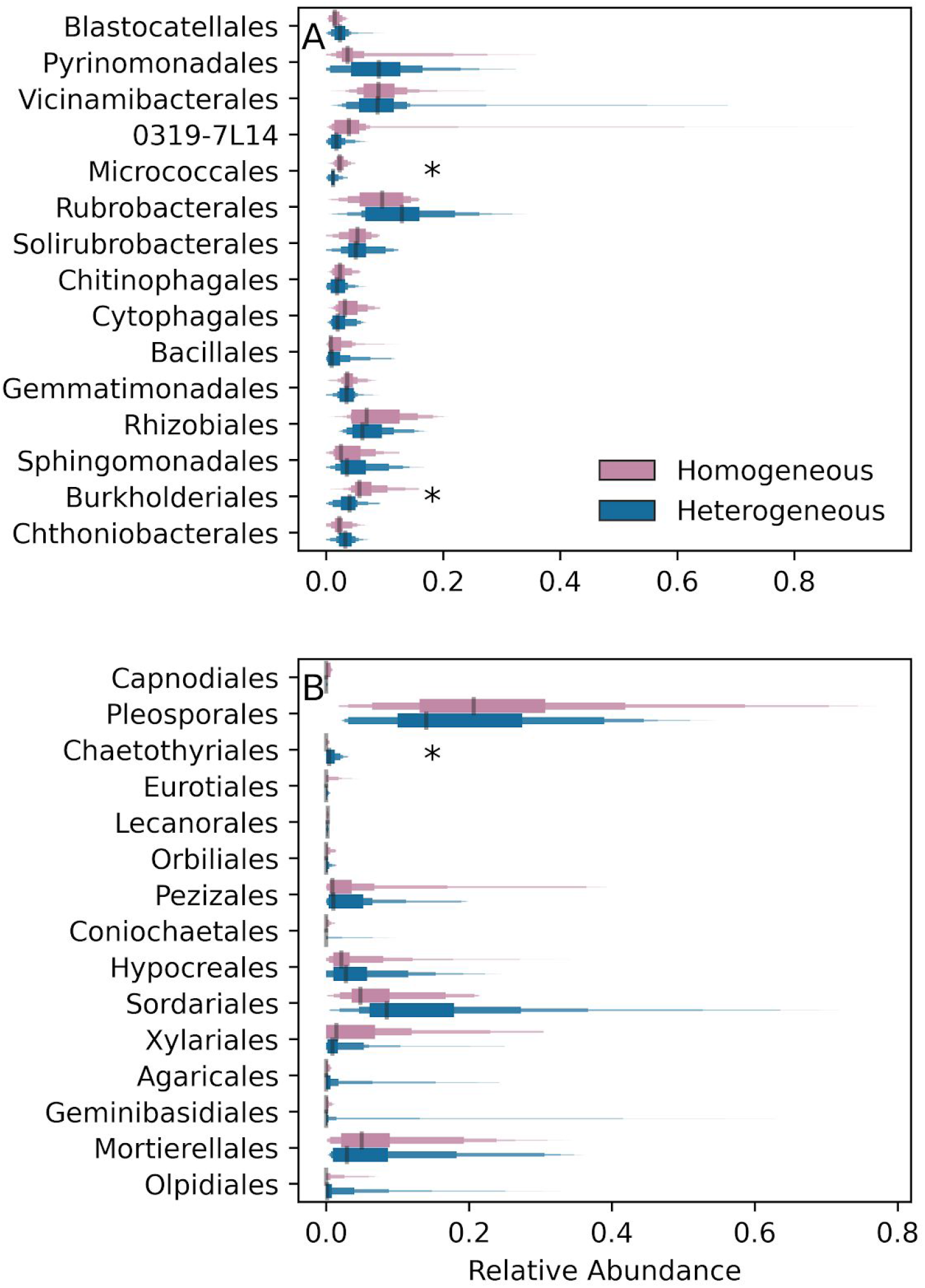
Order level relative abundance distributions in prokaryotes (A) and fungi (B). * – Significant difference between homogeneous and heterogeneous hillslopes according to Mann Whitney U-test

### 3.1 Microbial diversity

For biodiversity analyses we rarefied all the samples to 3000 reads post-filtration, following the guidance of rarefaction curves (Fig. S1), which indicated that rare taxa were sufficiently represented. The Faith-PD metric (Faith, 1992), accounting for the number of amplicon sequence variants (ASVs), their relative proportions and their phylogenetic relationships, was used to describe alpha diversity. According to a factorial ANOVA accounting for the geodiversity levels (heterogeneous vs. homogeneous) and the niche (shrubby patch vs. inter-shrub), neither factor had significant effects on prokaryotes or fungi.

Unlike alpha diversity, beta diversity, expressed as pairwise UniFrac distances (Lozupone and Knight, 2005), was affected by hillslope type. In a principal coordinates analysis (PCoA; Legendre and Legendre, 2012) based on the bacterial weighted UniFrac matrix (Fig. 1B), geodiversity classes appeared to be partially segregated along the second axis, which explained 10.7% of the total variance. The beta diversity of fungi was also partially explained by hillslope type, based on the first axis of an unweighted UniFrac PCoA analysis (Fig. 1C, 9.3% of the variance).

To formally test whether the hillslope type significantly explained beta diversity, we carried out a factorial ANOVA test, in which the within-homogeneous hillslopes pairwise UniFrac distances, the within-heterogeneous hillslopes pairwise UniFrac distances and between hillslope-type distances were considered as levels. Only distances between pairs of samples belonging to the same niche were considered. For prokaryotes, differences among the unweighted UniFrac distance distributions (Fig. 1D) were significantly explained by the hillslope type (p-value < 0.0067), a pattern which was marginally more pronounced in inter-shrub spaces than in shrubby patches (p-value = 0.02). Distances were smaller within the homogeneous hillslopes than within the heterogeneous hillslopes (q-value < 0.0061), and marginally smaller within the homogeneous hillslopes than between the hillslope type (q-value < 0.024), indicating that the homogeneous hillslopes might be more selective or restrictive for prokaryotes. Weighted pairwise distances were not explained by hillslope type in prokaryotes, indicating that the observed pattern was due to differences in taxa and not due to differences in their relative abundances.

In fungi, hillslope type significantly explained pairwise unweighted distances as well (p-value < 1.5 x 10^-9^), without difference between the niches (shrub patches vs. inter-patch spaces; p-value = 0.9). For fungi, it was the heterogeneous hillslopes that had the smallest pairwise distances (Fig. 1E; q-value < 0.0036 and q-value < 1.3 x 10^-9^), but homogeneous hillslopes pairwise-distances were also smaller than between hillslope-type distances (q-value < 0.004). Weighted pairwise distances were less significantly explained by hillslope type (p-value = 0.005) than the unweighted distances (p-value < 1.5 x 10^-9^), indicating again that the difference in the taxa between the heterogeneous and homogeneous hillslopes was more important than the differences in their relative abundances.

We similarly tested the niche effect on beta diversity, considering only within hillslope-type distances. For prokaryotes, weighted pairwise distances within the niches were smaller than between the niches (p-value < 5 x 10-6; Fig. 1F), an effect evident in both homogeneous and heterogeneous hillslopes (p-value = 0.2). The effect was smaller when considering unweighted distances (p-value < 0.007), indicating that changes in relative abundances rather than in the taxa themselves were the key difference between the niches. Interestingly, niche differences generated negligible effects for fungi (Fig. 1G; p-value = 0.2 for weighted-UniFrac and p-value = 0.05 for unweighted UniFrac distances).

### 3.2 Explanatory ASVs – total variance

Explanatory taxa were investigated with BiPlot analyses (Legendre and Legendre, 2012) to identify the taxa explaining the total variance and with logistic regression classifications (Yu et al., 2011) and the ANCOM procedure (Mandal et al., 2015) to identify the taxa that diverged between the homogeneous and heterogeneous hillslopes. The most important bacterial ASVs for the total variance, following the importance definition of Legendre and Legendre (2012; Fig. 1B – biplot arrows) were *Acidobacterium* (ASV 71e3), *Arenimicrobium luteum* (ASV 8ea5), Vicinamibacteraceae sp. (ASV 6d1f) and *Rubrobacter* sp. (ASV dfb4). These ASVs aligned with the second PCoA axis (Fig. 1B). Thus, *Acidobacterium* (ASV 71e3) and *Arenimicrobium luteum* (ASV 8ea5) were usually more abundant in the heterogeneous hillslopes while Vicinamibacteraceae sp. (ASV 6d1f) and *Rubrobacter* sp. (ASV dfb4) were usually more abundant in the homogeneous hillslopes, but were obviously affected by factors other than geodiversity or niche. For fungi, the most important taxa explaining the total variance (Fig. 1C, biplot arrows) were three *Geminibasidium* sp. ASVs (649c, 6fcf and 0ae1) and one *Stagonosporopsis* sp. ASV (49da), but their abundance was not shaped by the hillslope type or niche differences.

### 3.3 Explanatory ASVs and prokaryotic genes – homogeneous vs. heterogeneous hillslopes

According to a logistic regression, prokaryote relative abundances were well explained by the hillslope type (Fig. 3A; mode *R^2^* = 0.89). Out of the 20 taxa that best explained the regression (Fig. 3B), ASVs which were best associated with a homogeneous-hillslope classification included Vicinamibacteraceae sp. (691f; order Vicinamibacterales), *Arthrobacter* sp. (fe74; Micrococcales), Bacteroidetes sp. (fdc0; Cytophagales), Nitrosomonadaceae – NMD1 (b2fa; Burkholderiales), Nitrosococcaceae sp. – wb1-P19 (a360; Nitrosococcales), Chitinophagaceae sp. (bfa2; Chitinophagales), Burkholderiaceae sp. (e5df; Burkholderiales), *Gemmatimonas* sp. (24b7; Gemmatimonadales), Nitrososphaeraceae sp. (cd0b, 1505; Nitrososphaerales), Acidimicrobiia-IMCC26256 sp. (6946), *Adhaeribacter* sp. (c3a1; Cytophagales), *Microvirga* sp. (87e5; Rhizobiales), *Chthoniobacter* sp. (de3f; Chthoniobacterales), Actinobacteria – 0319-7L14 (40e8) and Solirubrobacterales – 67-14 (5fe9). The higher relative abundance of Vicinamibacteraceae sp. (691f) in homogeneous hillslopes was also supported by the ANCOM test. Many of these taxa are cellobiose degrading aerobes (Huber and Overmann, 2019a, 2019b, 2018; Janssen, 2015; Jones and Keddie, 2006; Kämpfer et al., 2011; Reichenbach, 1992; Zhang et al., 2019), whose increase may cause a shift in the soil’s redox properties, as cellobiose is a reducing sugar (Schellenberger et al., 2011). Ammonia oxidizers (Holmes et al., 2001; Prosser et al., 2014) and iron oxidizers (Hu et al., 2018) are also a striking component of this cohort. Prokaryote taxa that were best associated with a heterogeneous hillslope classification were Pyrinomonadaceae sp. – RB41 (3351, 9fe6, 7254; Pyrinomonadales) and *Rubrobacter* sp. (1649; Rubrobacterales). Of these taxa, only *Rubrobacter* sp. degrades cellobiose (Albuquerque et al., 2014; Wüst et al., 2016).

**Fig. 3.**
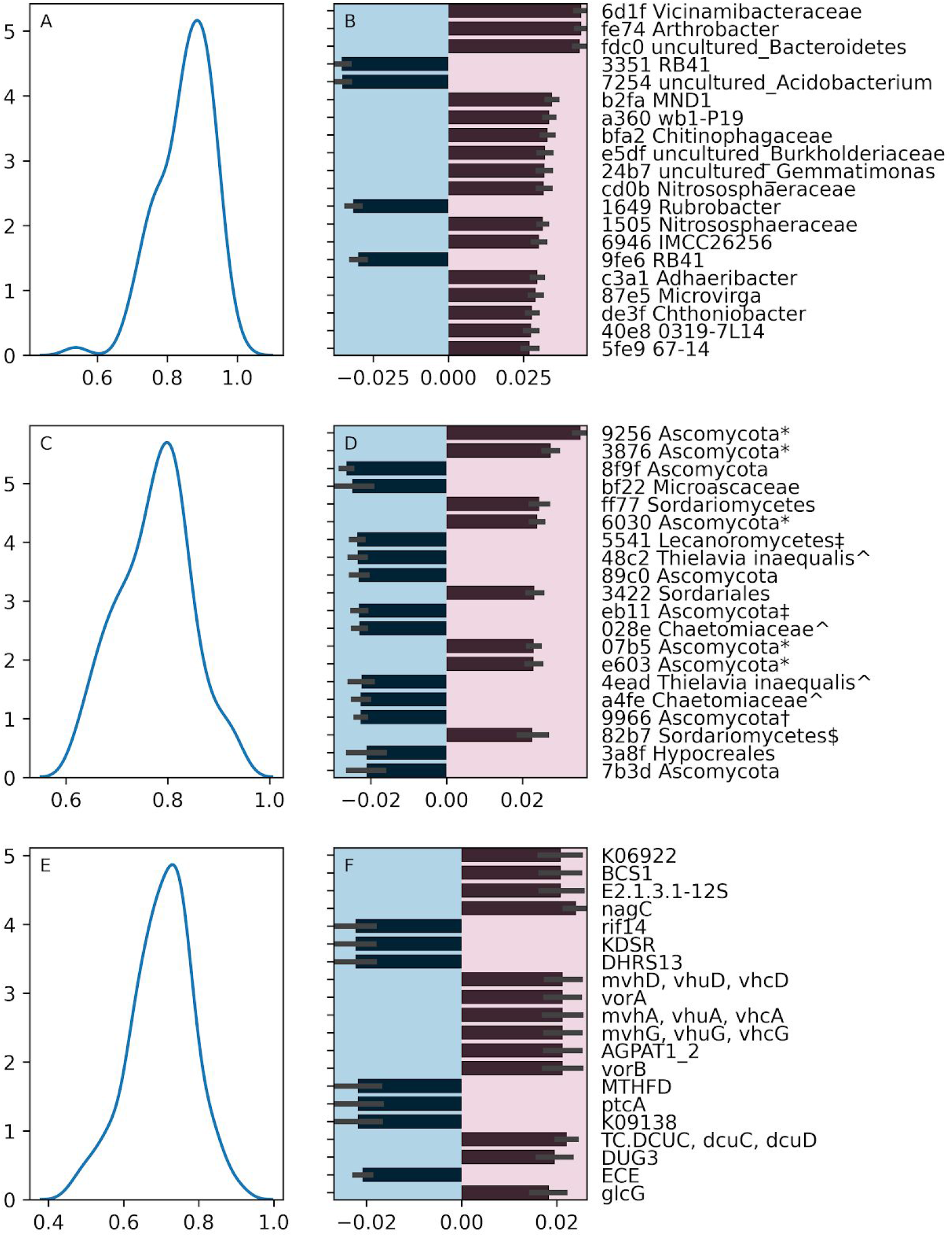
Explanatory taxa and bacterial genes. The distribution of *R*^2^ (A, C and E), and the coefficients with the largest absolute size (B, D, C) resulting from logistic regressions of bacterial relative abundances (A and B), fungal relative abundances (C and D) and bacterial genes (E and F) between the two hillslope types. A *R*^2^ distribution was generated in each regression by repeating the analysis for 100 randomly split training and testing datasets. In all iterations, geodiversity was equally stratified in the training and testing sets. Taxonomic and phylogenetic insights from Fig. S2: *-Monophyletic and assigned to Phaeosphaeriaceae. $-Hypocreales. ^-Monophyletic *Thielavia inaequalis.* †-Monophyletic Ascomycota. ‡-Monophyletic Lecanoromycetes.

A good fit to hillslope type was obtained for fungal relative abundances as well (Fig. 3C; mode *R^2^* = 0.8), with five Ascomycota ASVs (9256, 3876, 6030, 07b5, e603), which were monophyletic and assigsned to Phaeosphaeriaceae according to a phylogenetic analysis (Fig. S2), along with a Sordariomycetes sp. ASV (ff77), Sordariales sp. (3422; Sordariomycetes) and another Sordariomycetes ASV (82b7; Hypocreales sp. according to Fig. S2), were best associated with a homogeneous hillslope classification. The higher relative abundance of Phaeosphaeriaceae (Ascomycota ASV 9256) in homogeneous hillslopes was also supported by an ANCOM test. Phaeosphaeriaceae and Sordariomycetes are associated with plant pathogens and saprobes (Maharachchikumbura et al., 2015; Phookamsak et al., 2014). Conversely, three distinct Ascomycota sp. ASVs (8f9f, 89c0, 7b3d), Microascaceae sp. ASV (bf22; Microascales), two monophyletic Lecanoromycetes sp. ASVs (5541; eb11, Fig. S2), four *Thielavia inaequalis* ASVs (48c2, 028e, 4ead, a4fe; Sordariales, Fig. S2) and another Hypocreales sp. (3a8f), were best associated with a heterogeneous hillslope classification. An order level ANCOM test also supported the higher relative abundance of Chaetothyriales in heterogeneous hillslopes. Microascaceae (Sandoval-Denis et al., 2016), Chaetothyriales (Teixeira et al., 2017), Hypocreales (Maharachchikumbura et al., 2015) and *Thielavia inaequalis* (Srivastava et al., 1966) are associated with plant and/or vertebrate pathogens or decomposers, like fungi of the homogeneous hillslopes. However, Lecanoromycetes, which contain most of the lichen forming fungi (Miadlikowska et al., 2006), represent a very different functional group. To comment on possible artifacts in our ITS1 phylogenetic analysis (Fig. S2), monophyletic ASVs were manually inspected to confirm that the sequence divergence was genuine and not an ASV assembly artifact. Intraspecific divergence of ITS sequences in fungi was previously reported (Nilsson et al., 2008).

The fit of prokaryotic genes’ relative abundances, as reconstructed by PiCRUST2 (Langille et al., 2013), with the hillslope type, was lower than that of taxonomic relative abundances (Fig. 3E; mode *R^2^* = 0.72). Genes which were best associated with a homogeneous hillslope classification (Fig. 3F) included BCS1, a microbial 12S subunit protein, nagC, mvhD, vorA, mvhA, mvhG, AGPAT1_2, DCUC, DUG3 and glcG. They are involved in aerobic breathing and the tricarboxylic acid cycle (Hou et al., 2015; Janausch et al., 2002), disaccharide degradation (El Qaidi and Plumbridge, 2008), F420 oxidation (Stojanowic et al., 2003) and Ferredoxin activity (Charon et al., 1999). Conversely, rif14, KDSR, DHRS13, MTHFD, ptcA and ECE were best associated with a heterogeneous hillslope classification. The rif14 gene is a part of the Rifamycin B production pathway (Xu et al., 2003), DHRS13 is involved in the production of extracellular superoxides (Diaz et al., 2013), also promoted by the reduction of NADP+ (Diaz et al., 2013) by MTHFD (Röttig and Steinbüchel, 2013). KDSR is involved in sphingolipid metabolism in membranes of some anaerobes (Olsen and Jantzen, 2001). The gene ptcA is involved in putrescine metabolism and in one of the metabolic pathways producing gamma-aminobutyric acid (Chen et al., 2011; Jorge et al., 2017).

## 4. Discussion

In order to understand the functioning of ecosystems, it is important to account for the spatial structure of microbial communities. This structure is directly connected to spatial changes in turnover rates of many microbially mediated processes. More often than not, microbial communities and their functions are compared among different ecosystems (Nelson et al., 2016; Prober et al., 2015; Tedersoo et al., 2014; Wen et al., 2017), among distinctly different land uses, treatments or host classes (Espenberg et al., 2018; e.g., Giné et al., 2016; Krotman et al., 2020; Sher et al., 2012) or at a millimetric spatial scale, among minute ecological niches and pedology classes (Upton et al., 2019; Yergaliyev et al., 2020). Our study investigated the relationship between the structure of the soil microbial community and geodiversity at a hillslope scale, a few hundred square meters, within a single landform. Recent studies, which tied the hillslope-scale geodiversity variations with shrub mortality in this dryland region (Stavi et al., 2019, 2018a), illustrated that this scale of geodiversity may be an important determinant of microbial diversity.

In this study, we discovered that the hillslope type affected the structure of soil microbial communities. Particularly, the presence and absence of ASVs varied between the two hillslope types. For prokaryotes, the homogeneous hillslope samples were most similar to one another while for fungi, heterogeneous hillslopes were most similar to one another. This result may indicate that the magnitude of ecological drift (sensu Gilbert and Levine, 2017) was inverse for fungi and prokaryotes, as follows. Conditions were more restrictive to prokaryotic diversity in the homogeneous hillslopes and for fungi in the heterogeneous hillslopes, whereas the stochasticity of processes shaping the community was higher for prokaryotes in heterogeneous hillslopes and for fungi in the homogeneous hillslopes. Notably, albeit significant, the observed differences were small and we cannot attest to their dynamics.

Using logistic-regression analyses, we attempted to understand whether the increased prokaryotic diversity and reduced fungal diversity in the heterogeneous hillslopes are beneficial in terms of regulatory ecosystem services, particularly with regards to plants. A very consistent difference in the presence and absence of prokaryotic ASVs was observed between the two hillslope types (mode R^2^ = 0.89), which displayed a striking increase in cellobiose degrading aerobes in the homogeneous hillslopes, along with an increase in ammonia oxidizers. This suggests that the homogenous soil community is less favorable for plants because they harm the ammonia supply by actively oxidizing it and by shifting the redox potential in the soil. Various oxidative activities and disaccharide degradation were also observed among the bacterial genes associated with homogeneous hillslopes. The functional analysis also indicated that heterogeneous hillslope taxa were more active in the deterrence of pathogens, with antibiotics and superoxide production. Possibly, they were also more active in the production of gamma-aminobutyric acid (GABA). In plants, GABA is at the crossroad of several biotic and abiotic stress response mechanisms, including the production of succinic acid and lactic acid. Succinic acid, in turn, promotes energy production through the maintenance of the tricarboxylic acid cycle, often inhibited in stress conditions (Li et al., 2017; Shelp et al., 2017). An exogenous source of GABA was shown to increase the concentration of both endogenous GABA and succinic acid (Hijaz and Killiny, 2019). To summarize, it would seem that homogeneous-hillslope prokaryotic communities reduce ammonia availability and provide less pathogen response mechanisms, and are therefore detrimental to plants, in comparison with the communities of heterogeneous hillslopes.

Interestingly, the inverse pattern observed for fungi, wherein the fungal beta diversity was higher in the homogeneous hillslopes, could also be interpreted as being associated with the improved vigor of plants in the heterogeneous hillslopes. According to the logistic regression analysis of fungal relative abundances (mode R^2^ = 0.8), the increased diversity in homogeneous hillslopes can be attributed to plant pathogens and saprobes. The main functional difference between the two geodiversity classes that can be inferred from this analysis is the loss of Lecanoromycetes in the homogeneous hillslopes, which contain most lichenised fungi. Epilithic lichens play a role in the regulation of soil-water dynamics, as they intercept raindrops and allow slow release of the absorbed water (Zedda and Rambold, 2015). In water-limited habitats this could be crucial for the performance of plants.

Key questions that require further study relate to the soil-water threshold beyond which heterogeneous hillslopes can no longer maintain a supportive microbial community for plants, and the temporal stability of the observed patterns. Some insight can be gained from a study that compared soil microbial functions between live and dead shrub patches in the Sayeret Shaked LTER station, at the onset of shrub mortality (Sher et al., 2012). The researchers reported a higher ammonia oxidation potential under the canopies of dead *N. mucronata* shrubs than under live plants. According to Stavi et al. (2019, 2018a), *N. mucronata* mortality was spatially associated with low geodiversity. Therefore, there is an agreement between their observations and ours. Sher et al. (2012) also suggested that the lack of soil-water, which brought about aerobic conditions, could explain the increase in ammonia oxidation and reduced denitrification under drought conditions, which corresponds with our observations of increased abundances of ammonia oxidizing bacteria in the homogeneous, more aerated, hillslopes. Therefore, the comparison between Sher et al. (2012) and the current study provides some evidence for the persistence of microbial diversity patterns during the prolonged drought episode.

## 5. Conclusion

Hillslope geodiversity is an important determinant, not only of plant diversity but also of microbial diversity. Interestingly, our results suggest that the positive effect of high geodiversity on shrub vigor is not limited to direct actions of abiotic factors. Microbially conferred regulatory ecosystem services that facilitate shrub survival are also positively affected. Compared with the heterogeneous hillslopes, the deterioration of the shrub community in homogeneous hillslopes occurs not only due to the direct effect of reduced soil-water storage, but also through the shift in microbial community, caused by the same degraded soil-water conditions.

## CRediT authorship contribution statement

**A. Szitenberg**: Investigation, Formal analysis, Validation, Visualization, Writing – original draft, Funding acquisition, Resources. **R. Alexander-Shani**: Investigation, Resources. **Hezi Yizhak:** Conceptualization. **I. Stavi**: Conceptualization, Investigation, Writing – review & editing, Funding acquisition.

## Data statement

The authors confirm that the data supporting the findings of this study are available under BioProject PRJNA697940 and Zenodo DOI: 10.5281/zenodo.4480328.

## Acknowledgements

The authors thank Michelle Finzi for English language editing of the manuscript. This publication is based upon work from COST Action G-Bike (CA18134), supported by COST (European Cooperation in Science and Technology).

## Appendix. Supplementary data

Fig. S1: 16S rRNA and ITS1 rarefaction curves https://doi.org/10.6084/m9.figshare.13669100

Fig. S2: ITS1 ASV phylogeny. ASV sequences have red labels and reference sequences (black) are from the UNITE database. Bullets at node bases represent a 65% bootstrap support or higher. https://doi.org/10.6084/m9.figshare.13669106

